# Latent disease similarities and therapeutic repurposing possibilities uncovered by multi-modal generative topic modeling of human diseases

**DOI:** 10.1101/2021.05.18.444550

**Authors:** Satoshi Kozawa, Hirona Yokoyama, Kyoji Urayama, Kengo Tejima, Hotaka Doi, Thomas N. Sato

## Abstract

Human diseases are characterized by multiple features such as their pathophysiological, molecular, and genetic changes. The rapid expansion of such multi-modal disease-omics space provides an opportunity to re-classify diverse human diseases and uncover their latent molecular similarities, which could be exploited to repurpose a therapeutic-target for one disease to another. Herein, we probe this underexplored space by soft-clustering 6,955 human diseases by multi-modal generative topic-modeling. Focusing on chronic kidney disease and myocardial infarction, two most life-threatening diseases, unveiled are their previously underrecognized molecular similarities to neoplasia and mental/neurological-disorders, and 69 repurposable therapeutic-targets for these diseases. Using an edit-distance based pathway-classifier, we also find molecular pathways by which these targets could elicit their clinical effects. Importantly, for the 17 targets, the evidence for their therapeutic usefulness is retrospectively found in the pre-clinical and clinical space, illustrating the effectiveness of the method, and suggesting its broader applications across diverse human diseases.

## Introduction

Human diseases are characterized by alterations in a multitude of features: genetics, molecular, cellular, inter-organ pathways, histopathology, physiology, microbiota, etc. Recently, this disease-omics data space is rapidly expanding and becoming readily available, enabling the comprehensive characterizations of diverse human diseases (Hasin et al., 2017; Perakakis et al., 2018; Reel et al., 2021).

For example, GWAS catalog (https://www.ebi.ac.uk/gwas/) and other similar databases provide a comprehensive list of genetic factors associated with thousands of human diseases and traits. KEGG (https://www.genome.jp/kegg/), Reactome (https://reactome.org) and other similar databases describe comprehensive molecular pathways. DisGeNET (https://www.disgenet.org) compiles altered gene expression, biomarkers, posttranslational modifications, genetic factors, drug-targets, etc. and their association with human diseases. Disbiome (https://disbiome.ugent.be/home) tabulates human disease-associated microbiota. There are also large numbers of drug-related databases (SIDER: http://sideeffects.embl.de, DrugBank: https://go.drugbank.com, FAERS: https://www.fda.gov/drugs/questions-and-answers-fdas-adverse-event-reporting-system-faers/fda-adverse-event-reporting-system-faers-public-dashboard, etc.) that comprehensively list therapeutic-indications, side-effects/adverse-events, targets, etc. of the drugs. Moreover, we can virtually identify cells and organs that express genes/proteins of interest at The Human Protein Atlas (https://www.proteinatlas.org), Human Cell Atlas (https://www.humancellatlas.org), and other similar open-resources.

Hence, this rapidly expanding multi-modal disease-omics data space provides an opportunity to re-classify diverse human diseases according to their multi-modal similarity metrics and to find previously underrecognized disease-disease similarities. Furthermore, such latent disease-disease similarities could be exploited to repurpose a therapeutic target for a disease to treat another.

The pioneering study built on the graph theory provided an overview of disease-disease similarities according to their single modality features, genetic variabilities (Goh et al., 2007). Since then, more sophisticated network-based and other approaches have evolved to characterize multi-modal nature of human diseases (Barabasi et al., 2011; Garcia Del Valle et al., 2021; Menche et al., 2015; Perakakis *et al.*, 2018; Reel *et al.*, 2021). Despite such tools and methods development, the ever-expanding multi-modal disease-omics space remains under-explored. Hence further in-depth probing of this data space is expected to uncover latent molecular mechanisms underlying non-classical under-recognized disease-disease similarities.

As an approach that could integrate multiple types of features to classify human diseases and measure their similarity metrics, topic modeling was brought to our attention. This algorithm has been applied to categorize social media information (Zheng et al., 2014), and also to image annotation and classification and computer vision (Roller and Schulte im Walde, 2013). This approach has also recently been applied to the classification of clinical notes (Wen et al., 2021) and RNA dual-omics (RNA, microRNA) data (Valle et al., 2022).

Based on these previous reports, we considered a use of the multi-modal topic modeling to re-classify diverse types of human diseases according to their multi-modal disease-omics features. Furthermore, such multi-modal disease classification could be exploited to identify highly similar diseases and to translate such relationship to the therapeutic targets-repurposing.

Hence, we set the following two-fold objectives in this study:

1. To map human disease similarities according to their multiple modalities of disease-omics features.
2. To infer latent disease-omics features as candidates for therapeutic-targets according to the above-described human disease similarity map.

Here we report a modified multi-modal generative topic modeling-based method that achieves these objectives and its effective performance. In addition, we also illustrate its applications to two globally most life-threatening human diseases, chronic kidney diseases and myocardial infarction.

## Results

### Therapeutic targets-mining and the inference of disease similarities by multi-modal generative topic modeling

The overall design of our approach is described in Figure 1A. We performed soft-clustering of 6,955 human diseases by a multi-modal generative topic modeling according to the similarity metrics of their associated multiple disease-omics features. We also use this multi-modal generative topic modeling to infer latent disease-omics features as candidates for therapeutic-targes and/or molecular/genetic biomarkers for the corresponding disease(s). In the next step, we utilize comprehensive multi-modal biological-omics and disease-omics data in combination with an edit-distance based pathway classifier to examine their putative therapeutic pathways. Through this secondary screening, the final candidates with biologically and therapeutically sound potentials are selected. The presence/absence of the evidence for successful therapeutic outcomes in the real-world is retrospectively examined in the pre-clinical and clinical data space.

**Figure 1.**
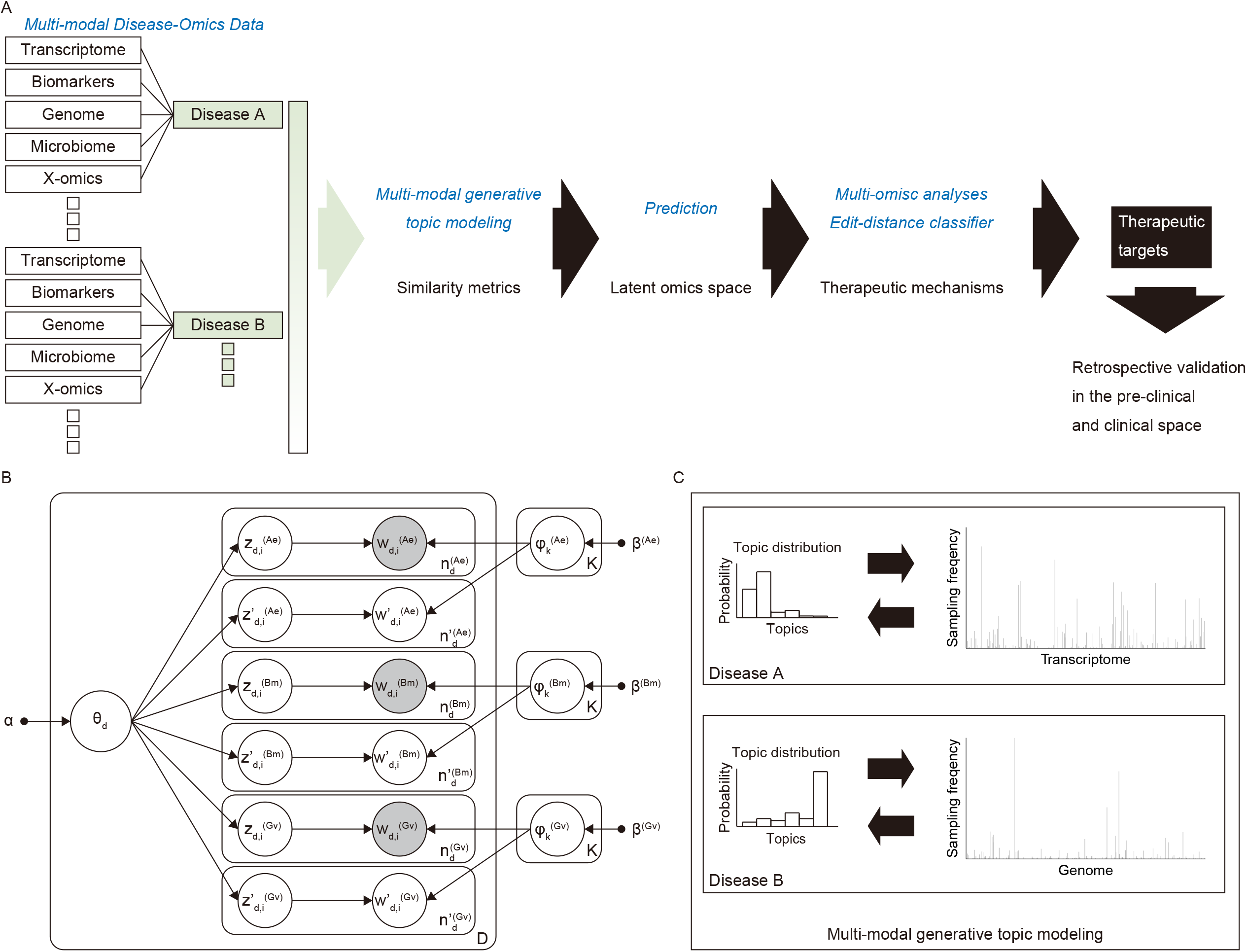
Schematic diagram of the multi-modal generative topic-modeling. (A) Overall scheme. Using the comprehensive multi-modal disease-omics datasets, the human diseases are soft-clustered by the multi-modal generative topic-modeling according to their multi-modal similarity metrics. The latent omics features are predicted as therapeutic target candidates. Their therapeutic mechanisms are deduced by utilizing multi-omics data and by the edit-distanced based classifier. These candidates represent repurposable therapeutic targets and their therapeutic utilities are further validated in the retrospective pre-clinical and clinical data space. (B) The multi-modal generative topic modeling algorithm. See “Multi-modal generative topic modeling of human diseases” of the Methods section for the details. (C) The predictions are based on the similarities of the probabilistic distributions of each human disease (Disease A, Disease B, etc.) across the topics. The predictions of the targets are output as sampling frequencies for each feature of the disease-omics modality (e.g., transcriptome, genome, etc.).

The multi-modal generative topic modeling that we employed is based on Latent Dirichlet Allocation (LDA) (Blei et al., 2003) (Figure 1B, also see “Multi-modal generative topic modeling of human diseases” in the Methods). We assume that each type of multi-modal disease-omics features exhibit the same probabilistic distribution pattern across the topics for the same disease. Hence, the latent and/or missing modality feature space of a disease is inferred according to the probabilistic distribution patterns of the existing features of the same and other diseases that are present in the training datasets (Figure 1C, also see “Multi-modal generative topic modeling of human diseases” in the Methods).

To evaluate the prediction performance of the method, we used a following set of multi-modal disease-omics features for 6,955 diseases: altered gene expression (Ae), biomarkers (Bm) and Genetic Variations (Gv) from the public multi-modal disease omics database, DisGeNET (https://www.disgenet.org). The prediction performance-validation is conducted by randomly dividing them to the 6,665 diseases as training data and the remaining 290 diseases as test data encompassing all 6,955 diseases. For each set of the test diseases, we removed their single modality features as prediction targets. Hence, the training disease-dataset consists of Ae, Bm, and Gv (AeBmGv), and the test disease-dataset consists of Ae and Bm (AeBm), or Bm and Gv (BmGv), or Ae and Gv (AeGv). We tested one modality at a time for each set of the test diseases. The prediction performance for the removed (i.e., missing in the test data) modality features in the test data is measured by Area Under receiver operating characteristic Curve (AUC) (Figure 2A, Table S1, also see “Calculation of AUC scores” in the Methods). The results show that virtually all AUCs for all diseases and all features are >0.8, supporting the effectiveness of the method.

**Figure 2.**
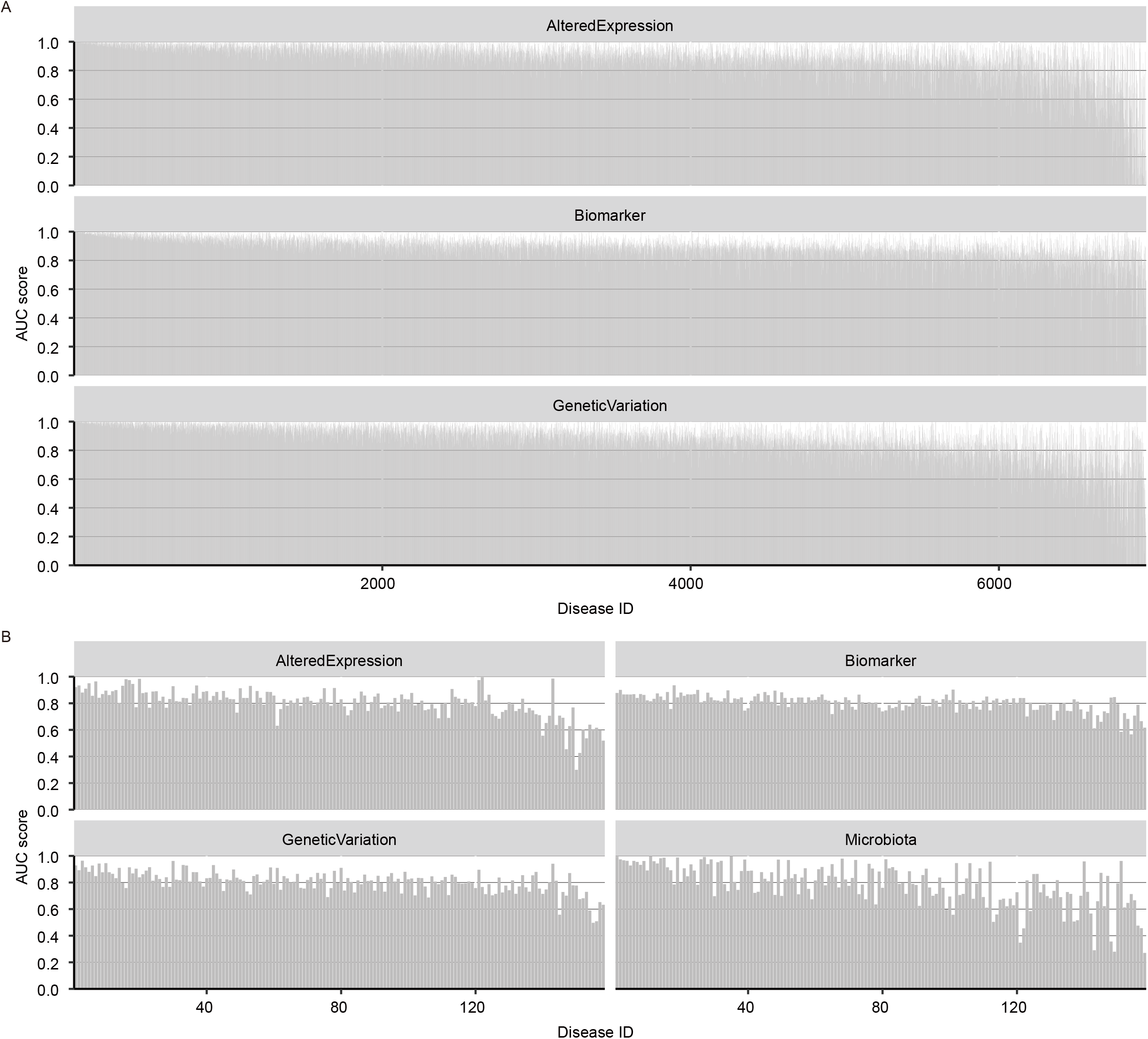
The AUC scores in the cross validations. (A) The AUC scores for the AlteredExpression, Biomarker, GeneticVariation modalities across 6,955 human diseases. The raw data are available as Table S1. (B) The AUC scores for the AlteredExpression, Biomarker, GeneticVariation, Microbiota modalities across 158 human diseases. The raw data are available as Table S2.

To test the modality-scalability of the method, we added another disease-microbiota dataset from the public database, Disbiome (https://disbiome.ugent.be/home) to the above AeBmGv datasets. This combination of datasets enabled us to soft-cluster 158 diseases all of which are annotated with the Ae, Bm, Gv, and microbiota (Mb) features. The performance is evaluated by leaving-one-modality-of-a-disease-out cross validation and by computing the AUC for each modality (i.e., Ae, Bm, Gv, Mb) of each disease (Figure 2B, Table S2, also see “Calculation of AUC scores” in the Methods). The result shows that AUCs are mostly >0.8 across all 158 diseases, supporting the modality-scalability of the method.

Deeper evaluation was performed for two globally most life-threatening diseases, chronic kidney disease (CKD) and myocardial infarction (MI). For this purpose, we use the above described AeBmGv datasets encompassing 6,955 diseases. To infer repurposable Ae, Bm, Gv from the other diseases as therapeutic-targets for CKD and/or MI, we purposely removed Ae, Bm or Gv features from CKD or MI and predicted the missing features. This approach yielded a list of predicted Ae, Bm, Gv features for each disease with all AUCs >0.8 (Figure 3, Tables S3 – S9).

**Figure 3.**
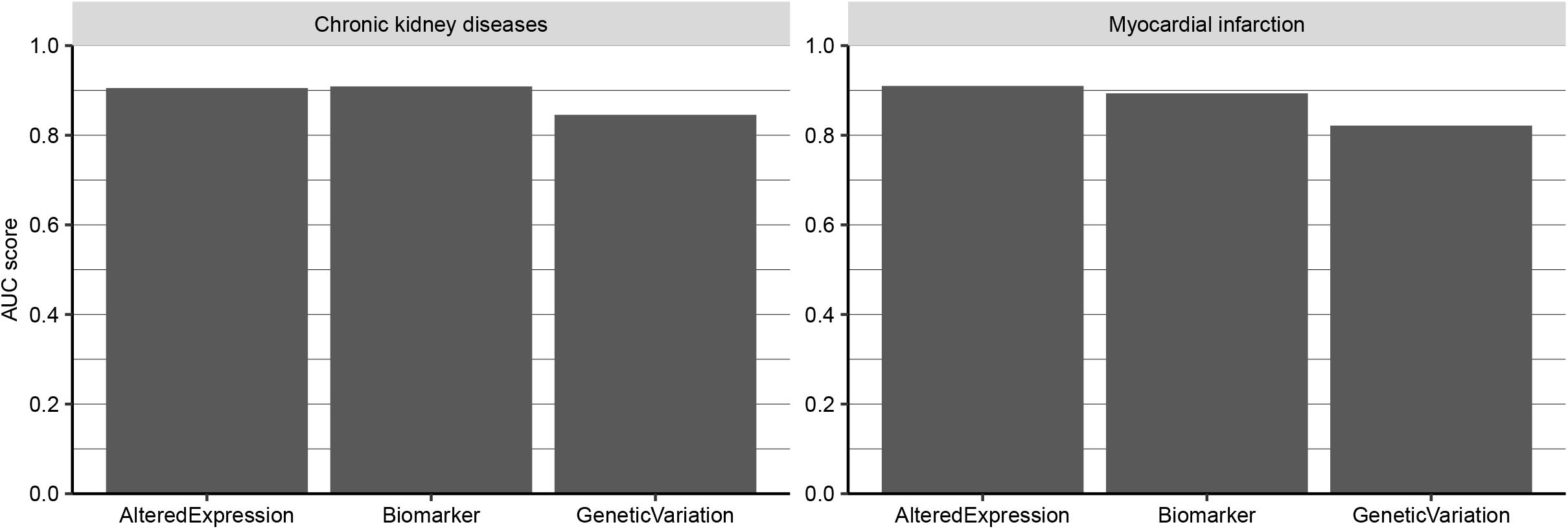
The AUC scores of each modality for the chronic kidney disease and the myocardial infarction. The AUC scores are calculated for AlteredExpression, Biomarker, GeneticVariation modalities for each disease (chronic kidney disease, myocardial infarction) and shown as bar graphs. The bars are shown as the AUC scores of 10x leave-one-modality/disease-out cross-validations. The raw data are available as Tables S3, S4, S5, S6, S7, S8, S9, S10, S11, S12, S13, S14, S15, S16, S17, S18.

From this list, the features that are present in the training datasets (i.e., those correctly predicted) were removed, leaving those that are absent in the CKD or MI data. Next, we used Youden’s index to select those that are considered as “statistically positive” by this criterion (Table S10, see also “Youden’s index” in the Methods). Through these selections, left are features that are absent in the training datasets for the corresponding diseases and regarded as statistically significant (Tables S11 – S16). Further selection was conducted by removing those that are labeled with other kidney/renal and cardiac/heart/cardiovascular related diseases (e.g., renal failure, heart failure, etc.), as we (i.e., without machine-learning methods such as ours in this paper) could easily postulate their repurposabilities to CKD and/or MI therapeutics. Through this additional selection step, we obtained a list of 30 and 57 molecular therapeutic candidates for CKD and MI, respectively, out of which 18 are shared by the two (Tables S17&S18). These candidates are particularly enriched in Ae, Bm and/or Gv of neoplasia (e.g., neoplasms, malignant neoplasms, neoplasm metastasis, malignant neoplasm of breast, primary malignant neoplasm, liver carcinoma, etc.) and mental/neurological disorders (e.g., schizophrenia, seizures, epilepsy, intellectual disability, etc.) (Figure 4, Table S19), unveiling their molecular similarities to the renal and/or cardiovascular diseases such as CKD and MI. Considering the relatively high AUCs of this inference method (Figure 3), this possibility is further supported.

**Figure 4.**
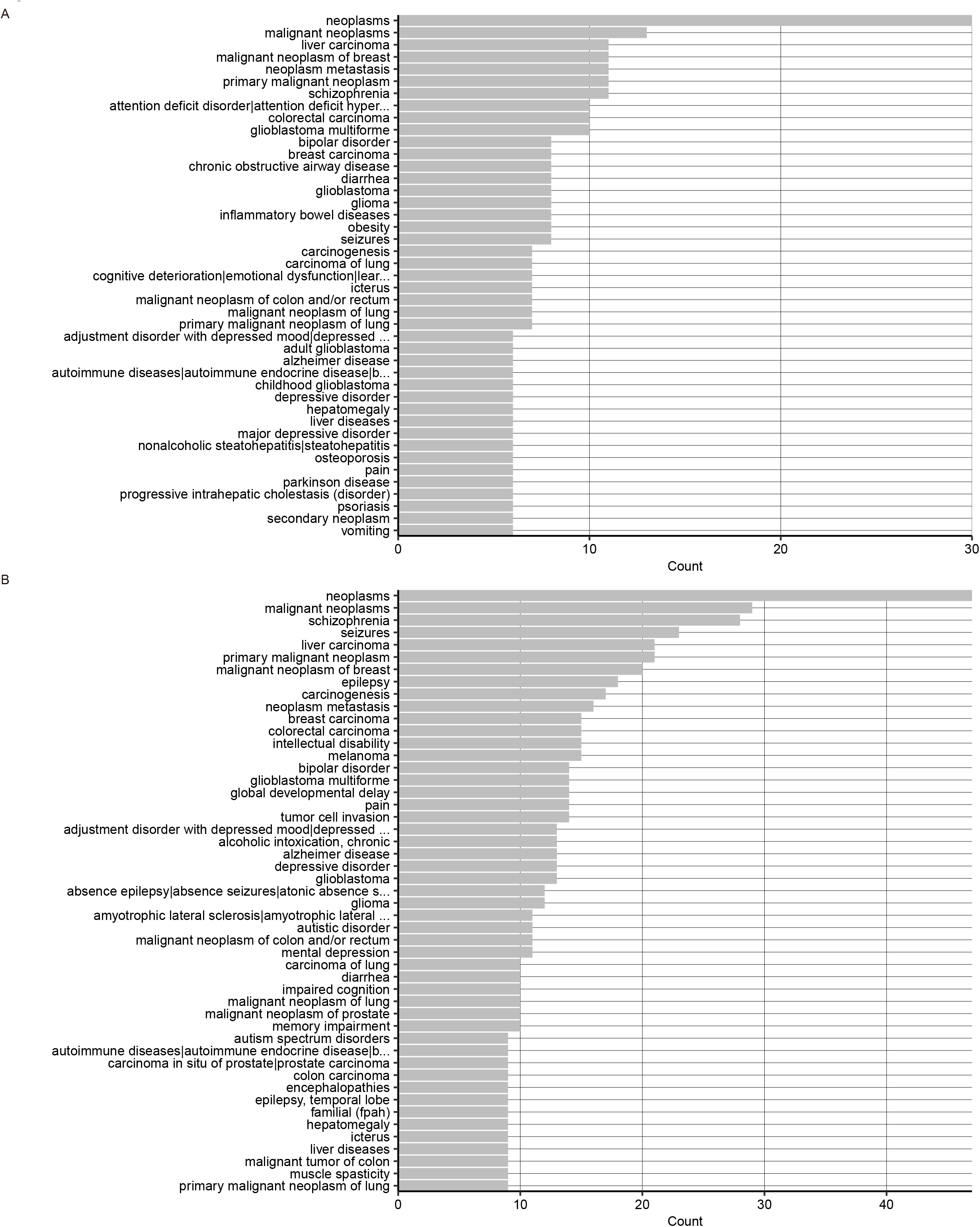
Human diseases determined as highly similar to the chronic kidney disease and the myocardial infarction by the multi-modal generative topic-modeling. (A) The diseases for the chronic kidney diseases. They are in the descending order of the multi-modal similarity metrics (Count). (B) The diseases for the myocardial infarction. They are in the descending order of the multi-modal similarity metrics (Count). The long disease names are shown as the first words and “..”. The raw data are available as Table S19.

### Therapeutic mechanisms of the inferred targets

To gain mechanistic insights into the putative therapeutic actions of the predicted target molecules, we identified their expression patterns in the human body using a comprehensive human protein/gene expression database (Figure 5, see also “Organ/cell expression enrichment analysis” in the Methods). At the organ-level, the CKD and MI targets are enriched in the liver and the brain, respectively (Figure 5A, B, Tables S20&S21). At the single cell-level, while cautions in the interpretations are necessary due to the apparent biases in the cell-type representations in the human single-cell transcriptome databases, we observe some enrichment in the hepatocytes and bipolar cells for CKD and MI, respectively (Figure 5C, D, Tables S22&S23). The result suggests that these organs/cells may serve as therapeutic targets for the respective diseases.

**Figure 5.**
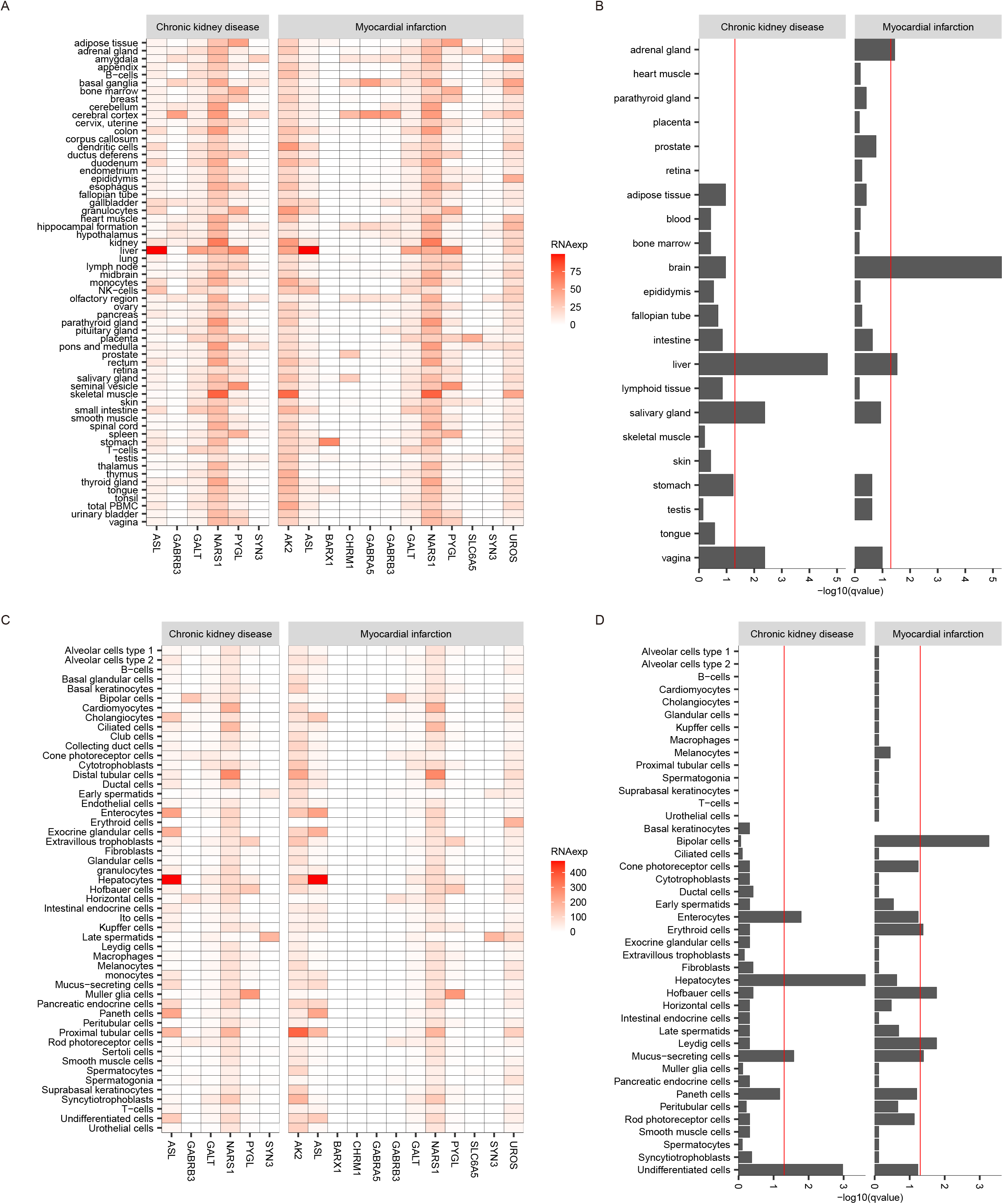
The organ/cell expression patterns of the predicted therapeutic targets for the chronic kidney disease and the myocardial infarction. (A) The heatmap indicating the expression levels for each predicted target (bottom) in each organ (left) for each disease (top). The raw data are available as Table S20. (B) The organ enrichment analysis result for the predicted targets for each disease (top). The enrichment levels are indicated as −log10(q-values). The raw data are available as Table S21. (C) The heatmap indicating the expression levels for each predicted target (bottom) in each cell-type (left) for each disease (top). The raw data are available as Table S22. (D) The cell-type enrichment analysis result for the predicted targets for each disease (top). The enrichment levels are indicated as −log10(q-values). The raw data are available as Table S23.

Further mechanistic insights were gained by the enrichment analyses of biological pathways and functions using KEGG and GO databases (Figure 6, Tables S24 – S28, see also “GO enrichment analysis” and “KEGG enrichment analysis” in the Methods). The analyses found the enrichment of the MI targets in neural-pathways and -functions (Figure 6, Table S24). These analyses, together with the expression pattern results, suggest that the nervous system functions and pathways are potential therapeutic targets for MI. In contrast, no enrichments are found for the CKD targets, instead they are sparsely encompassed across multiple biological pathways and functions (Tables S25 - S28).

**Figure 6.**
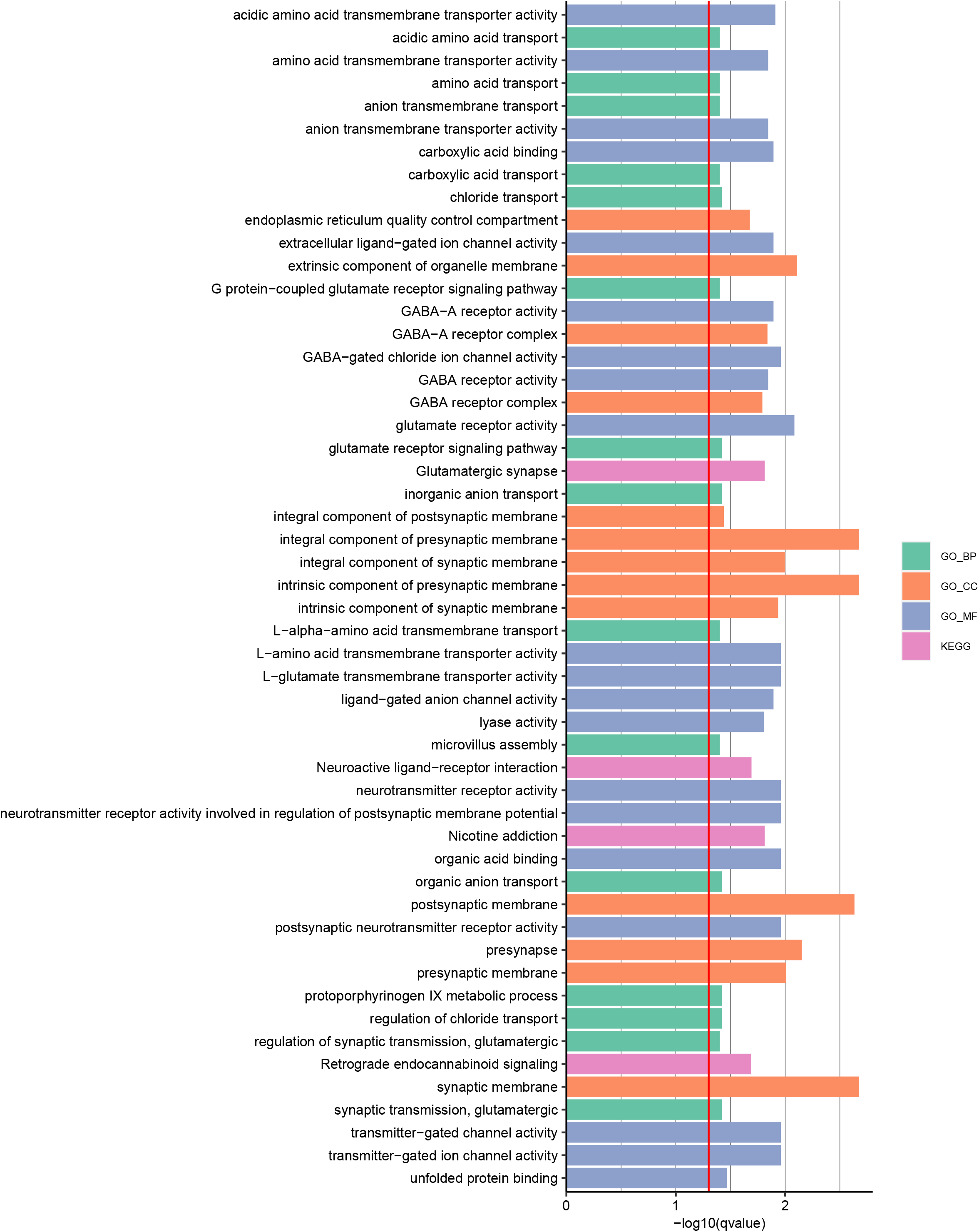
KEGG and GO enrichment of the predicted therapeutic targets for the myocardial infarction. The enrichments for each KEGG pathways and GO terms are shown as bar graphs. The enrichment is indicated as −log10(q-values). Those with −log(q-values) <0.05 are shown. The raw data are available as Tables S24, S25, S26, S27, S28.

Next, we designed an analysis to distinguish between two possible therapeutic mechanisms: 1) the predicted targets function via the therapeutic pathways that are also targeted by the clinically-approved treatments for CKD and/or MI, or 2) they function via other pathways that are distinct from those targeted by the clinically-approved treatments for the corresponding diseases. To address this objective, we measure similarity metrics of the biological and therapeutic pathways by a machine-learning classifier (Figure 7, see also “Edit-distance classification” in the Methods). This classifier uses the edit-distance, specifically Levenshtein distance, to measure the similarity metric between the pathways. In this method, the similarities of all possible pathways of the target candidates to each clinically-approved therapeutic pathway for each disease are computed (Figure 7, see also Methods). This is repeated for all pair-wise combinations for each disease and the computed edit-distances are used as input data for the corresponding disease-pathways classifier (Figure 7, see also “Edit-distance classifier” in the Methods). We applied this method to the CKD and MI classifiers to determine the extent to which the possible candidate molecule-driven pathways and the clinically-approved therapeutic pathways for CKD and/or MI share their pathways each other. The holdout validation shows that this method is highly reliable as indicated by the high prediction performance measures (i.e., accuracy scores>0.94, precision scores>0.71, recall scores>0.85, F1 scores>0.81) for both CKD and MI (Tables S29&S30).

**Figure 7.**
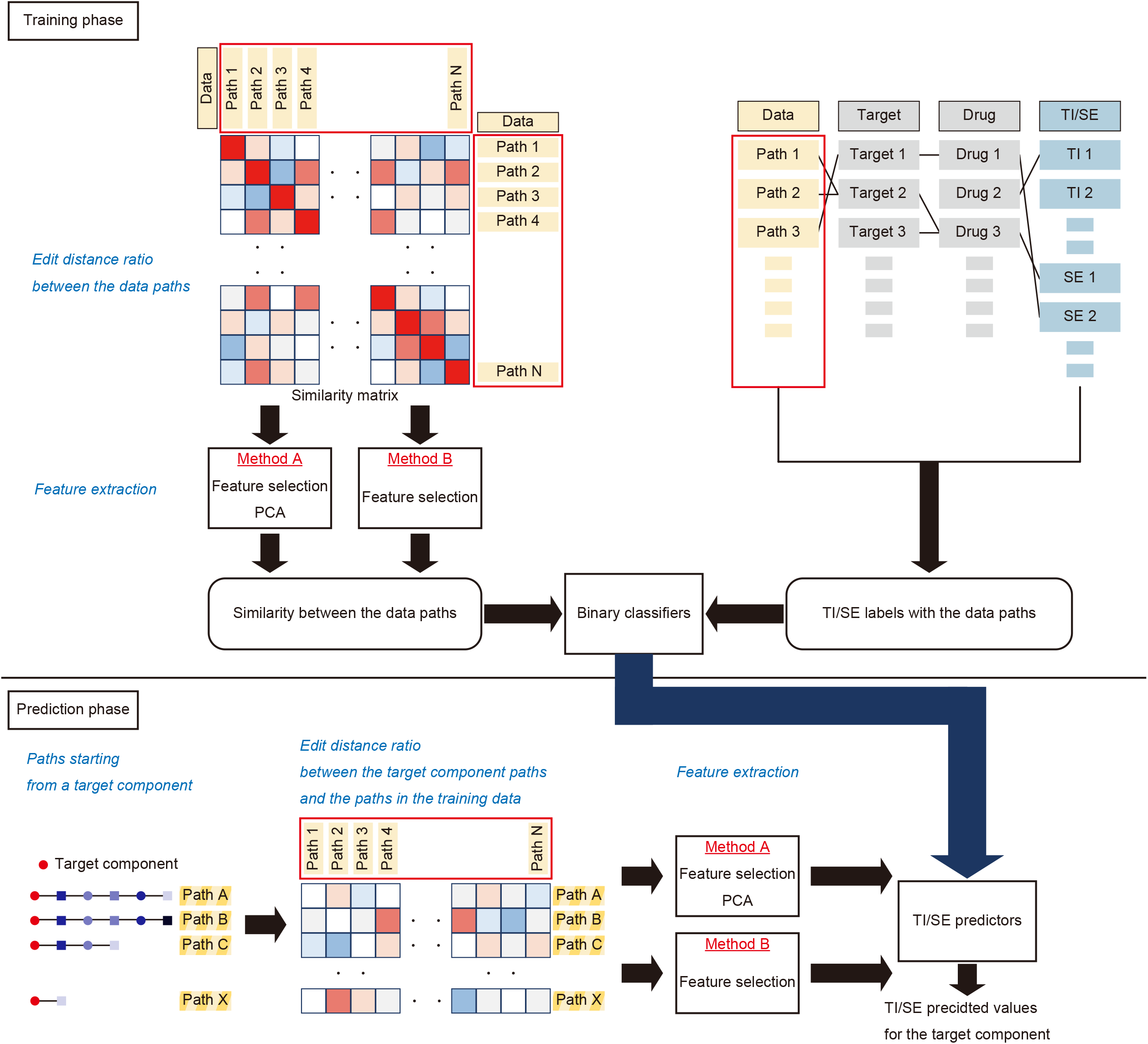
Schematic diagram of the edit-distance based classifier See “Edit-distance based classifier” in the Methods for the detailed step-by-step description. The raw data are available as Tables S29, S30.

This analysis found none of the CKD candidates share their pathways with the known CKD targets (Figure 8, Tables S31&S32). Moreover, none belong to the same KEGG pathways (Figure 8, Tables S31&S32). For MI, two candidates (ASL, LAMTOR1) are predicted as outside the known MI pathways, nor do they belong to the same KEGG pathways (Figure 8, Tables S33&S34). Additionally, nine other candidates (AK2, GABRA5, GABRB3, GALT, GRM7, PILRA, PRKG2, PYGL, GNPAT) are also predicted as outside the known MI pathways, although they belong to the same KEGG pathways as the MI pathways (Figure 8, Tables S33&S34). In contrast, two MI candidates (CHRM1, GRM3) and the known MI targets share the parts of their pathways (Figure 8, Tables S33&S34).

**Figure 8.**
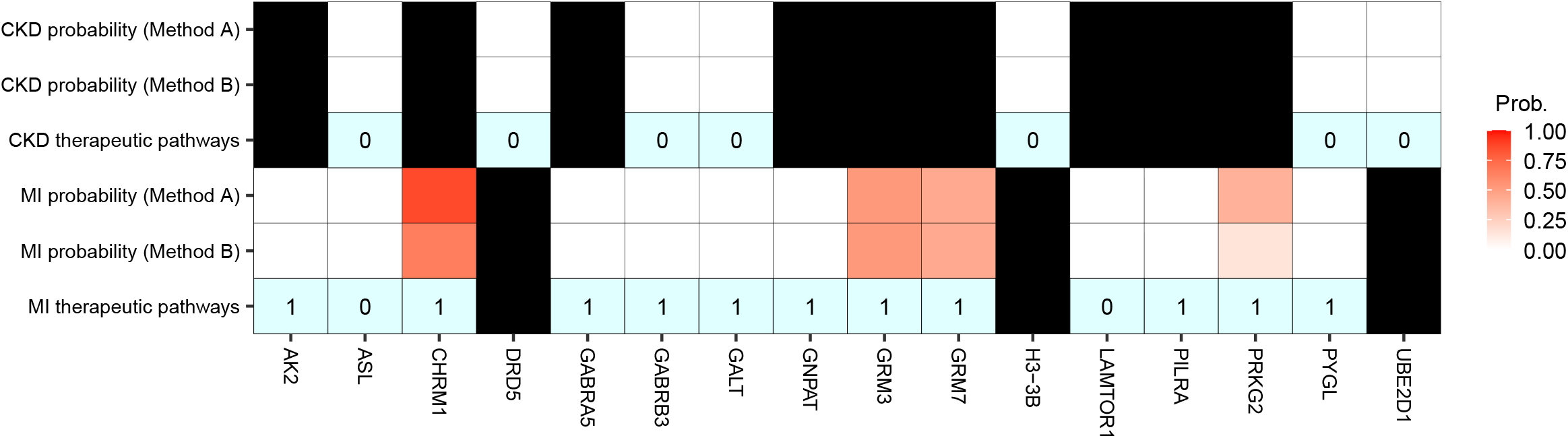
The deduced therapeutic pathways of the predicted targets. The probabilities that the targets elicit their predicted therapeutic effect(s) via the clinically approved therapeutic pathways for each target disease (CKD: chronic kidney disease, MI: myocardial infarction) is shown as heatmap (CKD/MI probability on the left). For which of the diseases (CKD vs. MI) each target are predicted is indicated by open (predicted for) or filled (not-predicted for) cells. Whether the targets are within or outside the same KEGG pathway(s) of the known targets of the clinically approved drugs for the corresponding target disease are indicated as “1” and “0”, respectively, in the corresponding cell (CKD/MI therapeutic pathways on the left). The raw data are available as Table S31, S32, S33, S34

Therapeutic-targeting elicits both favorable and unfavorable effects. The above analyses provide insights into the putative mechanisms of the former for the predicted targets. The latter is referred to as side-effect (SE). Hence, we applied the above-described edit-distance classier method to the prediction of SEs that could accompany the therapeutic-targeting of the candidate molecules (Figure 7, see also “Edit-distance classifier” in the Methods). In this study, we focused on the 176 serious adverse outcomes. The hold-out validation shows F1 scores>0.5 for 124 out of the 176 SEs (Table S35), suggesting that this prediction method is relatively useful. This prediction found four candidates (AK2, ASL, PILRA, PYGL) that are free of the selected 176 serious adverse outcomes (Figure 9, Tables S36&S37), suggesting that they are less harmful therapeutic-targets.

**Figure 9.**
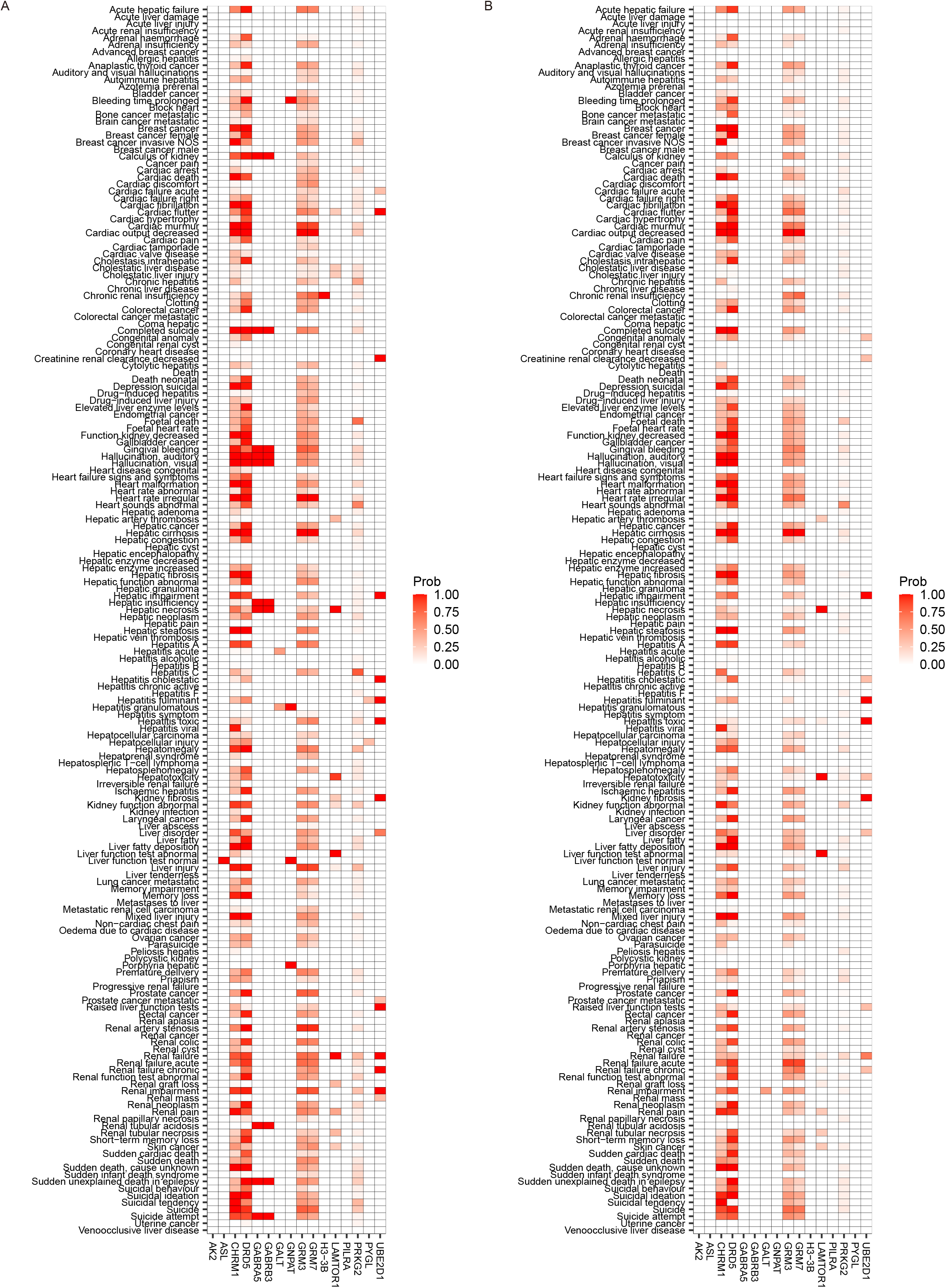
The side-effects inferred by the edit-distance based classifier for the predicted therapeutic targets. (A) The probabilities for the 176 serious side-effects (left) are indicated as the heatmap for the CKD targets (bottom). The raw data are available as Tables S35, S36. (B) The probabilities for the 176 serious side-effects (left) are indicated as the heatmap for the MI targets (bottom). The raw data are available as Tables S35, S37.

### Retrospective pre-clinical and clinical validation of the 69 targets

Next, we retrospectively evaluated the predicted therapeutic-utilities of the 69 targets in the pre-clinical and clinical data space (Table 1, Table S38). For the pre-clinical data space, we probed publications that might have been missed in the training datasets of this study. For the clinical data space, we conducted comprehensive data-mining of the past and on-going clinical trials at the clinicaltrials.gov (https://www.clinicaltrials.gov). The result is summarized in Table 1, implicating the therapeutic usefulness of the 17 out of 69 targets for the corresponding or other renal and/or cardiovascular diseases.

For example, knockout mice for AK2, an MI candidate predicted to function via pathways distinct from the known MI pathways (Figure 8, Tables S33&S34), is reported to show cardiac dysfunctions (Zhang et al., 2021). This mouse study may suggest a role of AK2 in cardiac development and/or function, hence AK2 could serve as a therapeutic target for cardiovascular diseases such as MI.

Deficiency of ASL, a candidate for both CKD and MI predicted to function also outside the known MI pathways (Figure 8, Tables S33&S34), is a rare genetic disorder resulting in argininosuccinic aciduria, a defective urea cycle condition leading to the insufficient breakdown/removal of nitrogen from the body, and consequently the patients develop hypertension (Kho et al., 2018). Hypertension is a known risk factor for both renal and cardiovascular diseases including CKD and MI (Clemmer et al., 2022; Ku et al., 2019). A clinical trial (NCT02252770) was conducted to evaluate the benefit of a nitric oxide dietary supplementation to argininosuccinic aciduria patients, but no outcomes are yet reported.

A cross-transplantation studies using the kidneys from the DRD5 knockout mice, CKD candidate predicted to function outside the known CKD pathways (Figure 8, Tables S31&S32), results in hypertension and cardiac dysfunctions in this mouse model (Asico et al., 2011). Furthermore, both hypertension and cardiac dysfunctions are known risk factors for renal diseases such as CKD (Clemmer *et al.*, 2022; Ku *et al.*, 2019), hence supporting the prediction of DRD5 as a therapeutic candidate for CKD.

GUCY2C, guanylate cyclase 2C, is predicted as a therapeutic target for both CKD and MI (Table 1). In the clinical trial (NCT03217591), therapeutic effects of a soluble guanylate cyclase stimulator, IW-1973 (a.k.a., Praliciguat) for diabetic nephropathy/diabetic kidney diseases were evaluated. The outcomes show several metrics supporting further investigation of Praliciguat for diabetic kidney diseases (Hanrahan et al., 2020).

Unilateral ureteral-obstruction in mice results in the upregulation of H3-3B, a CKD candidate predicted to function via independent pathways from the known CKD pathways (Figure 8, Tables S31&S32), in the kidneys (Shindo et al., 2018). In addition, a knockdown experiment of histone cell cycle regulation defective homolog A (HIRA) in a normal rat kidney cell (NRK-52) causes the decreased H3-B3 expression and increased fibrogenesis (Shindo *et al.*, 2018). Furthermore, in patients with IgA nephropathy, H3-3B immune-stains positively correlate with kidney fibrosis (Shindo *et al.*, 2018). These results support a therapeutic candidacy of H3-3B for renal diseases including CKD.

Genetic mutation of PYGL, a candidate for both CKD and MI predicted to function outside the known CKD or MI pathways (Figure 8, Tables S31 - S34), in human prevents effective glycogen breakdown in the liver leading to glycogen storage diseases (Zhan et al., 2021). While no clinical implications for CKD or MI or other renal/cardiovascular diseases in the patients of these conditions are recorded (NCT02385162), our expression analysis (Figure 4) suggests that the liver is a potential therapeutic target for both CKD and MI. Hence, PYGL could serve as a therapeutic target for CKD, MI and/or other renal/cardiovascular diseases.

UBE2D1, a CKD candidate predicted to function outside the known CKD pathways (Figure 8, Tables S31&S32), is inferred as a potential biomarker for diabetes-related sepsis by a machine-learning pipeline using public databases (Wang et al., 2022). Diabetes is a known risk factor for CKD (Shahbazian and Rezaii, 2013), hence, further supporting the candidacy of UBE2D1 for a CKD target as deduced herein.

These and other evidence summarized in Table 1 further supports the therapeutic possibilities of the candidates reported in this study for CKD and/or MI and/or other renal and/or cardiovascular diseases.

## Discussion

In this paper, we applied a multi-modal soft-clustering to the multiple disease-omics datasets and uncovers latent molecular similarities across 6,955 human diseases (Figures 1 - 4). By exploiting these molecular similarities, we identified 69 targets that could be therapeutically repurposed for CKD and/or MI treatments (Table 1). The comprehensive omics analyses, in combination with an edit-distance based classifier, found their underlying therapeutic mechanisms (Figures 5, 6, 8, 9). Importantly, we found the evidence retrospectively supporting the predicted therapeutic utilities of the 17 targets in the pre-clinical and clinical data space (Table 1).

Here, we used the soft-clustering of human diseases by multi-modal generative topic modeling, enabling the detection of subtle differences in the multi-modal features of the diseases (Figure 1). As a result, we were able to develop an algorithm that exhibits mostly AUC>0.8 for predicting the missing modality features of 6,955 human diseases (Figure 2A). In this study, we tested this method with three modalities, Ae, Bm, Gv, for 6,955 diseases (Figure 2A), and four modalities, Ae, Bm, Gv, Mb, for 158 diseases (Figure 2B). The result shows both sets result in virtually equivalent performance, suggesting the scalability of the method with additional modalities.

This method identifies the molecular features shared by CKD/MI and non-renal/non-cardiovascular diseases such as neoplasia and mental/neurological disorders (Figure 4), indicating a latent underlying mechanism shared among these diseases. The neoplasia can be regarded as a partial cellular reprogramming, as it is accompanied by the aberrant activations of large number of genes (Buganim et al., 2012; Suva et al., 2013; Ward and Thompson, 2012; Xing et al., 2020). This phenomenon could be reflected on the molecular similarities between the renal/cardiovascular diseases (e.g., CKD and MI) and the neoplasia. It is also recently reported that MI accelerates breast cancer via innate immune reprogramming (Koelwyn et al., 2020). This clinical observation might be a consequence of their molecular and mechanistic similarities as predicted in this study. Moreover, various clinical observations also suggest that CKD and cancer are mutual risk-factors, but without any clear molecular mechanisms (Wong et al., 2016). Hence, it is possible that the predicted molecular mechanisms/pathways shared by CKD and neoplasia reported in this paper may be an underlying molecular mechanism of these clinical observations.

Virtually all peripheral organs such as the liver, the kidney, the heart, etc. are under the control of neural inputs and these organs feedback their physiological information to the neural organ such as the brain (Imai et al., 2008; Underwood, 2021). Hence, such inter-organ neural feed-forward and feed-back loops could be reflected on the similar molecular features and underlying mechanisms of the renal/cardiovascular diseases (e.g., CKD and MI) and the mental/neurological disorders as predicted by the method reported herein. In support of this possibility, many mental disorders are prevalent in CKD patients (Simoes et al., 2019). Furthermore, myocardial infarction is often followed by deteriorated mental health conditions (De Hert et al., 2018; Lloyd, 1987). Despite such clinical evidence, no concrete molecular mechanisms explaining these clinical observations remain unknown. Thus, the common molecular mechanisms and/or pathways described in this study could be the ones.

In this study, such disease-disease similarities are further translated to the repurposing of therapeutic targets across the diseases, identifying 69 molecules that could be therapeutically targeted for CKD and/or MI treatments (Table 1). Their expression patterns and KEGG/GO analyses indicate they are enriched in the brain and the metabolic organs such as the liver and their physiological functions (Figures 5&6). These results are coherent with the molecular similarities between the CKD/MI and mental/neurological disorders described in this paper. They are also consistent with the fact that many of the renal/cardiovascular diseases including CKD and MI are broadly regarded as metabolic and life-style diseases (Sharifi-Rad et al., 2020; Thomas et al., 2011).

The edit-distance based classifier shows two types of therapeutic mechanisms by which these 69 candidates could elicit their effects in the treatments of CKD, MI and/or other renal-/cardiovascular-diseases. Those that function via the pathways that are also targeted by the drugs approved for the corresponding diseases (i.e., CKD, MI), and the others that function independent from them (Figure 8). The independent pathways may be a part of the previously unknown molecular mechanisms underlying the corresponding disease(s). In this case, their therapeutic-targeting could lead to the development of “first-in-class” drugs for the corresponding diseases. In contrast, those within the already-targeted pathways are activated or inhibited by the existing drugs, hence they could be further developed by adding new indications for the diseases that are described as molecularly similar in this paper.

The edit-distance based classifier is also applied to evaluate putative SEs that could accompany the therapeutic targeting of these candidate molecules (Figure 9). The result shows the four (AK2, ASL, PILRA, PYGL) are less harmful targets. This analysis provides a useful information for selecting out those that are likely less toxic prior to spending labor-, time-, and cost-intensive pre-clinical and clinical studies during the therapeutic development.

The likeliness of the repurposabilities of the predicted CKD and MI targets is further strengthened by the retrospective finding of the therapeutic implications of the 17 targets in the pre-clinical and clinical-trials data space, despite their absence in the training datasets (Table 1). This retrospective validation, together with the high AUC scores obtained by the cross-validation (Figures 2&3) and the modality scalabiliy (Figure 2), demonstrates the effectiveness of the method in uncovering latent disease-disease similarities and therapeutic repurposing possibilities across diverse diseases and modalities. Hence, the method is expected to be effective, not only for CKD or MI, but also for other types of diseases and with different and/or additional combinations of disease feature modalities.

## Acknowledgements

We thank K. Sugisaka, R. Takahashi, R. Kitaura, R. Ishikawa for administrative assistance. We are also grateful to the members of Sato lab at ATR and Karydo TherapeutiX, Inc. for advice and discussion throughout the course of this work. This work was supported in part by Innovative Science and Technology Initiative for Security Grant Number JPJ004596 ATLA Japan (T.N.S.), JST ERATO Grant Number JPMJER1303 Japan (T.N.S), Nakatani Foundation (T.N.S) and AMED under Grant Number JP21he2102002 (T.N.S).

## Author contributions

T.N.S. conceived the idea of the project, designed the study and supervised the overall research project. S.K. designed and performed the multi-modal generative topic modeling analyses. S.K. and H.Y. designed and performed the edit-distance based classifier. S.K., H.Y., K.U., K.T., H.D. established the datasets. T.N.S., S.K., H.Y., K.U., K.T. wrote the manuscript.

## Declaration of interests

All authors are employees of Karydo TherapeutiX, Inc.

Correspondence and requests for materials should be addressed to T.N.S.

## Data and Code Availability

The code reported in this paper is available at: https://github.com/skozawa170301ktx/MultiModalDiseaseModeling

## Methods

### Multi-modal generative topic modeling of human diseases

The generative topic modeling is based on Latent Dirichlet Allocation (LDA) (Blei *et al.*, 2003), enabling to perform soft-clustering of human diseases based on their multi-modal disease-omics features. This method can also predict latent and/or missing disease-omics features from the naïve model (Figure 1). The generative model is described as bellow. Let 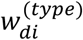 be the i-th component of disease *d* acquired from dataset that indicated by (type), 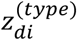 be the topic number of 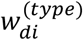, *θ_d_* be the topic probability of disease *d*, and 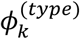 be the occurrence probability of component *v* of topic *k* at (type) dataset. In this paper, (type) is one of AlteredExpression (Ae), Biomarker (Bm), GeneticVariation (Gv), Microbiota (Mb). The topic number K was determined by community detection method (see “Community detection” section below). The generative model of this topic models was defined as follows:

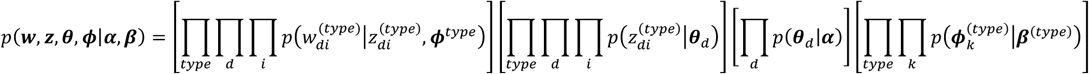

The exact formula of the elements of the joint distribution is described as:

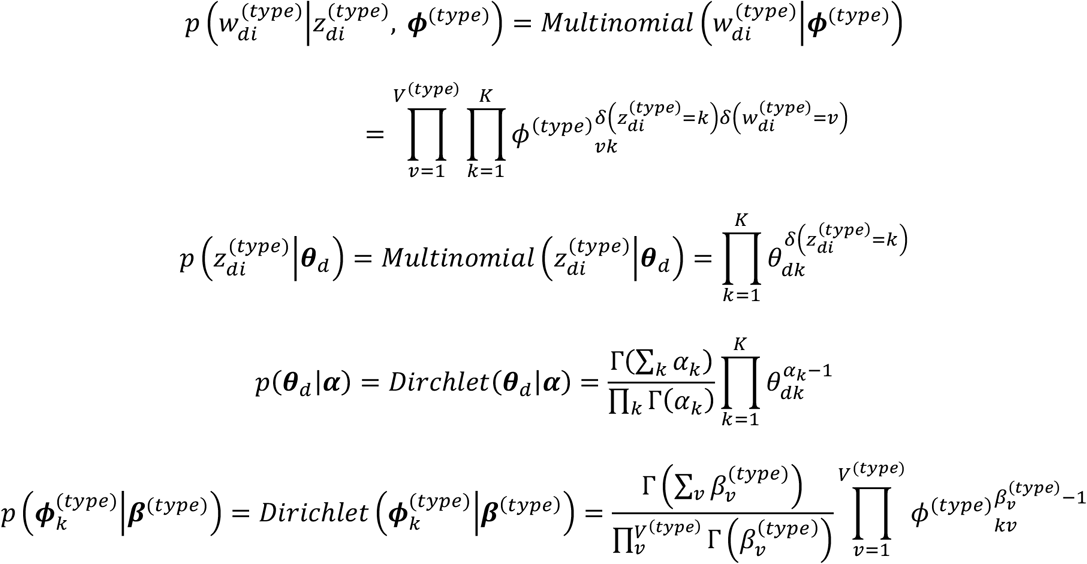

where, 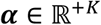 and 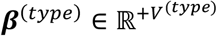 are the hyper parameters and we set these to a vector that elements have 0.1, *V*^(*type*)^ is the total kind of component at a (type) dataset, Γ(·) is the Gamma function and *δ*(·) is the Kronecker delta function. Based on the model, we estimated the posterior distributions of the variables 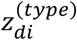, ***θ**_d_* and 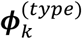 and also estimated a part of 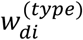 (it will be named 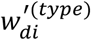) that not observed at some (type) dataset by using Gibbs sampling method. From the generative model, the conditional distributions of the variable were calculated as follows:

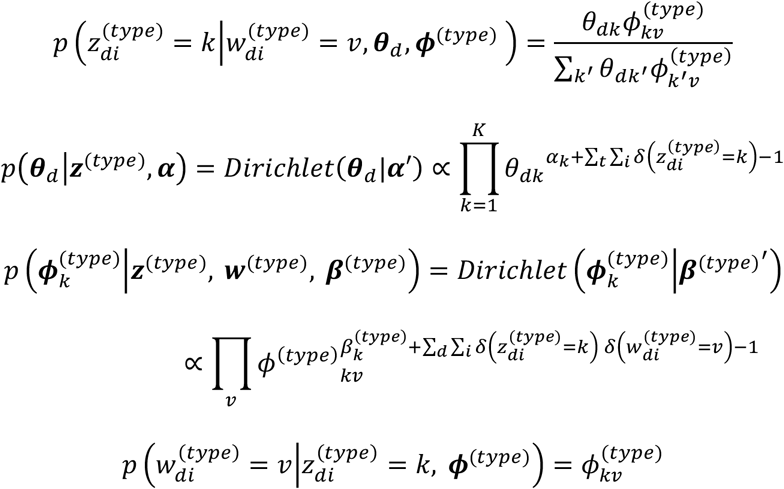

From these conditional distributions, we sample 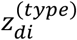 and 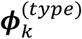 and 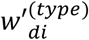, until its values converges. In this paper the number of iterations was set to 5,000, that was decided by manually decision. After the end of sampling, we can estimate the values of each variable by averaging the sampled values from the conditional distributions. The initial values of the variables are set as follows: The initial values of ***θ**_d_* and 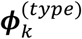 were set to the uniform distribution. The initial value of 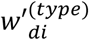 was set to the component that is random sampled from 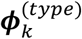. The total number of 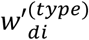 at disease *d* was decided by sampling the binominal distribution. The parameters for the binominal distribution were estimated by the maximum likelihood estimation method using the observed data. The estimated value of ***θ**_d_* represents the probability of topics at disease *d*. The likeliness of the missing values of disease *d* at (type) dataset can be inferred by sampling frequency of 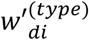.

### Community detection

The number of topics K was selected by a community detection method, the Louvain method. To apply this method, a undirected graph per dataset (Ae/Bm/Gv/Mb) was constructed. In the graph, if both of diseases have overlapping components, we allow an edge between the two diseases (i.e. nodes). The Louvain method was applied 20 times for each modality, and the most frequently obtained number of communities were selected for the community number for each modality. The maximum community number for each modality is selected as the K. The Louvain method is performed by Python package ‘python-louvain’.

### Calculation of AUC scores

To evaluate the likeliness of the missing component prediction, we calculated the AUC scores by using the result of the missing components prediction. The inputs are the Ae/Bm/Gv or Ae/Bm/Gv/Mb dataset where single modality components (i.e. Ae) were purposely left-out for a single disease. We then performed the multi-modal generative topic modeling as described in the previous Methods section on each of these input datasets, The likeliness of the missing components by the sampling frequency of 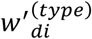. The frequency is defined as the prediction probability of the missing component 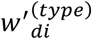, and the label can be defined whether the component is present or not in the original dataset, and the AUC scores can be calculated from the prediction probability and the labels at a disease in the (type) dataset. Calculation of the AUC scores were performed by function ‘roc_auc_score()’ in package ‘scikit-learn’.

### Youden’s index

To determine the cut-off threshold for the sampling frequency of 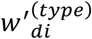, we used Youden’s index (Youden, 1950). Youden’s index is a ROC curve-based thresholding method. The thresholds for each modality for each disease was computed as follows: 1) True positive rate (TPR) and false positive rate (FPR) were computed using the function ‘roc_curve’ in the package ‘scikit-learn’, 2) The Youden’s index was then calculated by the formula, TPR – FPR for each threshold, 3) For each modality of each disease, the threshold which corresponds to the maximum Youden’s index was selected as the cut-off threshold for the corresponding modality for each disease.

### Multi-modal disease-omics datasets

The Ae, Bm, Gv data are from DisGeNET v7.0. (https://www.disgenet.org/downloads). The drug-target datasets are from DrugBank (https://go.drugbank.com/). The Mb data are from Disbiome (version on 11^th^ of November 2020, https://disbiome.ugent.be/home) (Janssens et al., 2018). The drug-therapeutic indications/-side effects data are from SIDER4.1 (http://sideeffects.embl.de/download/). The KEGG (https://www.genome.jp/kegg/) and GO (http://geneontology.org/) datasets are from the corresponding links. THE HUMAN PROTEIN ATLAS v 21.1. (https://www.proteinatlas.org) is used for the human organ/cell expression analyses. The table is downloaded from “25. Data from the Human Protein Atlas in tab-separated format” in the “DOWNLODABLE DATA” page (https://www.proteinatlas.org/about/download).

### UMLS IDs assignment to the disease names

To combine disease names acquired from multiple data sources, we add UMLS IDs to the disease names. The UMLS ID annotation was performed by ‘UMLS_AUI.extract_terminology (“ICD10”)’ function from Python library PyMedTermino (version 0.3.3) (Lamy et al., 2015). Prior to performing this function, ‘’s’ was replaced by a blank space in disease names. Following assigning the UMLS ID annotations, the UMLS IDs were combined by string ‘|’ if these UMLS ID have the same disease names. For the disease names that this UMLS ID annotation method failed, the actual disease names in the datasets were used and only those with the exact matching names were combined.

### Organ/cell expression enrichment analysis

To find the specific organ/cell-expression patterns for the target genes, we used a organ/cell enrichment analysis. The enrichment analysis was performed by using chi-square test of independence to evaluate the statistical significance of the enriched expression in the specific organ(s)/cell(s) detected for the genes of interest. We performed the test by making the 2×2 contingency table consisting of the appearance frequency of the genes of interest and that of the genes of interest in the particular tissue/cell. This table was used as the input to perform the chi-square test of independence using the Python function ‘scipystats.chi2_contingency()’.

### GO enrichment analysis

To find the specific gene ontology terms for the target genes, we used GO enrichment analysis. GO enrichment analysis was performed by R function ‘enrichGO()’ in package *‘clusterProfiler’*. The symbol names of the genes were converted to Entrez IDs by inputting enrichGO() using R function ‘bitr()’ in the package ‘*clusterProfiler*’.

### KEGG enrichment analysis

To find the specific KEGG pathways for the target genes, we used KEGG enrichment analysis. KEGG enrichment analysis was performed by R function ‘enrichKEGG()’ in the package ‘*clusterProfiler*’. Genes were handled as Entrez IDs as described above.

### Edit-distance based classifier

The overall design of the method, consisting of multiple steps, is schematically shown in Figure 7. Each step is described below:

#### Path extraction for the training phase

Each path starts with a known drug target. The drug-targets are from DrugBank. The drugs used in the training are those that are labeled with therapeutic indications and side-effects in SIDER4.1. Paths are from extracted from the KEGG database using Python package ‘networkX’. The databases are as described in “Multi-modal disease-omics datasets” of the Methods section above.

#### Edit-distance similarity computation

The edit-distances, specifically Levenshtein distances, between the paths were computed as follows:

The similarity of Path N and Path M was calculated as following:

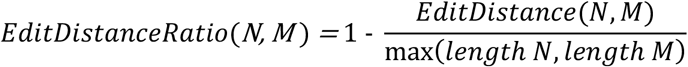

*Length X* is the number of elements in Path X. When two or more arguments are given to max(), it selects and returns the largest argument. Therefore, max(*length N, length M*) returns the largest number of elements between the Paths N and M. Then, the edit-distance was calculated by ‘*EditDistance*()’, using a dynamic programming algorithm (Navarro, 2001), as follows:

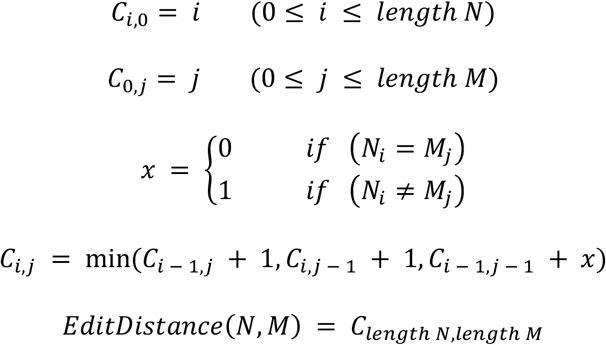

*N_i_* is the *i*-th element of paths N and *M_j_* is the, *j*-th element of path M. When more than one argument is given to ‘min()’, it selects the smallest argument. Finally, the computed edit-distance ratio between the paths N and M is used for their similarity metric.

#### Datasets for the edit-distance based binary classifier

The binary classifier of TIs and SEs was developed using LightGBM (Ke et al., 2017). We trained machine learning models by using the edit-distance based similarity metric between the paths, and their therapeutic-indication (TI) or side-effect (SE) labels. For the explanatory variables, we generated a symmetric matrix (hereinafter, referred to as A) with rows and columns as paths using the edit-distance ratios that are computed as above. The symmetric matrix A calculated above was input to the LightGBM as the explanatory variable following the feature selection step. The feature selection was performed in two different ways (Method A and Method B as below “Feature selection” and in Figure 7). For the response variables, the SEs or TIs for each drug target was obtained by combining drug-TI/SE (from SIDER, see the “multi-modal disease-omics datasets” section above) and drug-target (from DrugBank, see the “multimodal disease-omics datasets” section above). These TI and SE patterns for the drug-targets were input into the LightGBM as the response variables. The explanatory and response variables were divided to 9(training):1(test) for the holdout evaluation.

#### Feature selections

Feature selection for the explanatory variable was performed in two different ways (Method A and Method B) as follows:

For the method A, the paths with each of the known TIs or SEs were selected from the symmetric matrix A as the features for the corresponding TI and SE models. Then, principal component analysis (PCA) was performed on the matrix of each TI or SE to reduce the data dimensionality. The principal components were calculated until the cumulative contribution ratio reaches 0.99. The method B is without the PCA (Figure 7). Instead, the symmetric matrix A was divided on the drug target associated with the paths of the column, and within each divided matrix, the correlation matrix was calculated, and the column path was clustered using the group average method (i.e., a hierarchical method). The threshold was applied when the forming flat clusters was set to 0.2. We also selected the paths that are associated with each TI or SE as the features for the corresponding TI or SE models, and referred to them as A’. Based on the clusters of the paths as calculated above, this matrix A’ was sampled to reduce data dimensionality without PCA.

#### Training of the models

The dataset related to TIs or SEs and the dataset not related to it were downsampled at a ratio of 1:1 to train the model. Random under-sampling was performed on Python package ‘imblearn’. LightGBM was selected as machine learning algorithm. Bagging was done by creating 10 LightGBM models for each TI or SE. Hyperparameter tuning was performed by ‘optuna.integration.lightgbm’ (https://optuna.readthedocs.io/en/stable/reference/generated/optuna.integration.lightgbm.train.html). To verify the hyperparameter tuning, the training dataset was split in a ratio of 4:1. LightGBM was performed on Python package ‘optuna’.

#### Prediction of TIs and SEs for the feature components

The test path was generated as above by using a KEGG component as the starting node of the path. The edit-distances between this test path and all training paths for each TI or SE model was calculated as described in “Datasets for the predictions of therapeutic indications (TIs) and side-effects (SEs)” section above and used as input data. In the predictions, when the edit-distance results show ≥0.5 with 6 or more of the 10 models, the test path was determined as “true”, and when the results show otherwise, it was determined as “false”.

### The validation in the pre-clinical and clinical data space

The comprehensive pre-clinical and clinical data-mining was conducted via Google, PubMed and other literature search engines. For clinical data-mining, we also used the clinicaltrials.gov search engine (https://www.clinicaltrials.gov).

**Table 1.** The retrospective validation of the predicted therapeutic targets in the pre-clinical and clinical data space. The publications and clinical-trials are indicated as doi and clinicaltrial.gov NCT numbers, respectively. The raw data are available as Table S38.

| CKD | MI | Targets | Publication | ClinicalTrials.gov |
| --- | --- | --- | --- | --- |
|  | MI | ADRM1 |  |  |
| CKD | MI | AEBP1 | DOI:10.1186/s12967-021-03000-3 |  |
|  | MI | AK2 | DOI:10.1016/j.bbrc.2021.01.097 |  |
| CKD | MI | ASL |  | NCT02252770 |
| CKD | MI | ATP8B1 |  | NCT02094222 |
|  | MI | BARX1 |  |  |
| CKD | MI | BLOC1S2 | DOI:10.1038/cdd.2015.128 |  |
|  | MI | C15orf41 |  |  |
|  | MI | C19orf12 |  |  |
|  | MI | C2-AS1 |  |  |
|  | MI | CCDC115 |  |  |
|  | MI | CD79B |  | NCT04668365 |
|  | MI | CHRM1 |  |  |
| CKD |  | CRNKL1 |  |  |
| CKD |  | DRD5 | DOI:10.1681/ASN.2010050533 |  |
|  | MI | ECT |  |  |
|  | MI | EDS8 |  | NCT02165085 |
| CKD | MI | EPB41 |  |  |
|  | MI | ERLEC1 |  |  |
| CKD |  | ERVK-19 |  |  |
| CKD |  | FAM13A |  |  |
|  | MI | GABRA5 |  |  |
| CKD | MI | GABRB3 |  |  |
| CKD | MI | GALT |  | NCT02519504 |
|  | MI | GNPAT |  |  |
|  | MI | GRIK1 |  |  |
|  | MI | GRM3 |  |  |
|  | MI | GRM7 |  |  |
| CKD | MI | GUCY2C |  | NCT03217591 |
| CKD |  | H3-3B | DOI:10.1038/s41598-018-32518-8 |  |
|  | MI | HBB-LCR |  |  |
|  | MI | HBE1 |  |  |
|  | MI | HM13 |  |  |
| CKD | MI | HSD3B7 |  |  |
| CKD |  | HTR3A |  |  |
|  | MI | IKZF2 |  |  |
| CKD |  | IL36RN | DOI:10.1016/j.kint.2017.09.017 |  |
| CKD | MI | IREB2 |  |  |
|  | MI | LAMTOR1 |  |  |
| CKD |  | LINC01185 |  |  |
|  | MI | LMF1 |  | NCT03912181 |
| CKD | MI | NARS1 |  |  |
|  | MI | NDUFB2 |  |  |
| CKD |  | NME8 |  |  |
|  | MI | ODAM |  |  |
|  | MI | P3H3 |  |  |
|  | MI | PCDH19 |  |  |
|  | MI | PILRA |  |  |
|  | MI | PRKG2 |  |  |
| CKD | MI | PYGL |  | NCT02385162 |
|  | MI | RANBP2 |  |  |
|  | MI | RHAG |  |  |
|  | MI | SARM1 |  |  |
|  | MI | SLC17A7 |  |  |
| CKD | MI | SLC25A13 |  |  |
|  | MI | SLC6A5 |  |  |
| CKD | MI | SPAG8 |  |  |
|  | MI | SPAST |  |  |
| CKD | MI | SPATC1L | DOI:10.3892/mmr.2017.7334 |  |
|  | MI | SPNS1 |  |  |
|  | MI | STK26 |  |  |
| CKD | MI | SYN3 |  |  |
| CKD |  | THRSP | DOI:10.1155/2014/520281 |  |
| CKD | MI | TRP-AGG2-6 |  |  |
| CKD |  | UBE2D1 | DOI:10.1155/2022/9463717 |  |
| CKD | MI | UGGT1 |  |  |
|  | MI | UROS |  |  |
|  | MI | UST |  |  |
| CKD |  | ZDHHC13 |  |  |

## Supplemental information

**Figure S1.**
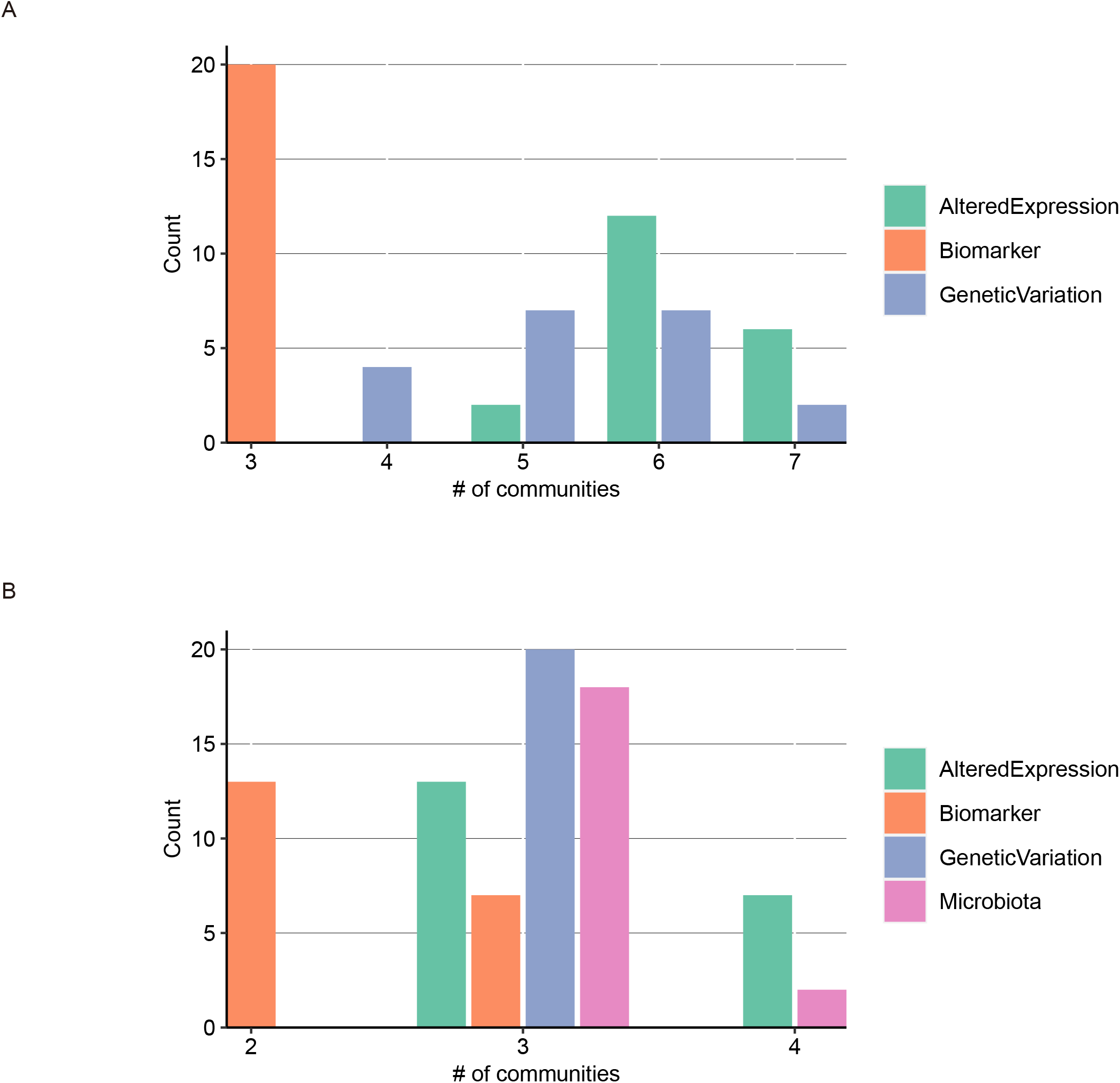
The community numbers, Related to Figure 1. (A) The numbers of communities found by the community detection method for AeBmGv datasets encompassing 6,955 diseases are shown as bar graph for each disease-omics modality. (B) The numbers of communities found by the community detection method for AeBmGvMb datasets encompassing 158 diseases are shown as bar graph for each disease-omics modality.

**Table S1.** The raw data of Figure 2A

Database: disease modality, key_c: UMLS ID of the corresponding disease – otherwise disease name, Disease_ID: disease number in the figure, Disease: disease name in the database (DisGeNET v. 7.0), AUC: AUC score.

**Table S2.** The raw data of Figure 2B

Database: disease modality, key_c: UMLS ID of the corresponding disease – otherwise disease name, Disease_ID: disease number in the figure, Disease: disease name in the database (Disbiome for Mb, DisGeNET v. 7.0 for the others), AUC: AUC score.

**Table S3.** The raw data of the AUC scores in Figure 3

element_name: disease-omics modality name, Disease: disease name, auc: AUC score.

**Table S4.** The list of predicted Ae features for CKD, Related to Figure 3.

The list of the predicted features (component) for Ae (AlteredExpression) modality for the chronic kidney disease (Disease). The count number (Count) and the frequency (Freq) in the prediction outputs are shown for each feature (component). The presence (1) and absence (0) of the predicted feature (component) in the training data are indicated in the “Obs” column.

**Table S5.** The list of predicted Bm features for CKD, Related to Figure 3.

The list of the predicted features (component) for Bm (Biomarker) modality for the chronic kidney disease (Disease). The count number (Count) and the frequency (Freq) in the prediction outputs are shown for each feature (component). The presence (1) and absence (0) of the predicted feature (component) in the training data are indicated in the “Obs” column.

**Table S6.** The list of predicted Gv features for CKD, Related to Figure 3.

The list of the predicted features (component) for Gv (GeneticVariation) modality for the chronic kidney disease (Disease). The count number (Count) and the frequency (Freq) in the prediction outputs are shown for each feature (component). The presence (1) and absence (0) of the predicted feature (component) in the training data are indicated in the “Obs” column.

**Table S7.** The list of predicted Ae features for MI, Related to Figure 3.

The list of the predicted features (component) for Ae (AlteredExpression) modality for the myocardial infarction (Disease). The count number (Count) and the frequency (Freq) in the prediction outputs are shown for each feature (component). The presence (1) and absence (0) of the predicted feature (component) in the training data are indicated in the “Obs” column.

**Table S8.** The list of predicted Bm features for MI, Related to Figure 3.

The list of the predicted features (component) for Bm (Biomarker) modality for the myocardial infarction (Disease). The count number (Count) and the frequency (Freq) in the prediction outputs are shown for each feature (component). The presence (1) and absence (0) of the predicted feature (component) in the training data are indicated in the “Obs” column.

**Table S9.** The list of predicted Gv features for MI, Related to Figure 3.

The list of the predicted features (component) for Gv (GeneticVariation) modality for the myocardial infarction (Disease). The count number (Count) and the frequency (Freq) in the prediction outputs are shown for each feature (component). The presence (1) and absence (0) of the predicted feature (component) in the training data are indicated in the “Obs” column.

**Table S10.** The Youden’s index for each modality feature, Related to Figure 3.

Database: modality, Disease: disease name.

**Table S11.** The list of predicted Ae features above the corresponding Youden’s index for CKD following the removal of those in the training data, Related to Figure 3.

Database: modality, key_c: UMLS ID of the corresponding disease (i.e., the chronic kidney disease), Disease: the chronic kidney disease, component: the name of the predicted feature, keg component id: KEGG ID number for the predicted feature (component), obs: 0 as they are absent in the training data, sample count: count number in the output, freq: frequency in the output, roc_cutoff: the threshold cut-off value based on the Youden’s index, DisGeNET Database: the modality in the DisGeNET database (i.e., the training data) where the corresponding feature (component) appears, DisGeNET Disease: the disease in the DisGeNET database (i.e., the training data) where the corresponding feature (component) appears.

**Table S12.** The list of predicted Bm features above the corresponding Youden’s index for CKD following the removal of those in the training data, Related to Figure 3.

Database: modality, key_c: UMLS ID of the corresponding disease (i.e., the chronic kidney disease), Disease: the chronic kidney disease, component: the name of the predicted feature, keg component id: KEGG ID number for the predicted feature (component), obs: 0 as they are absent in the training data, sample count: count number in the output, freq: frequency in the output, roc_cutoff: the threshold cut-off value based on the Youden’s index, DisGeNET Database: the modality in the DisGeNET database (i.e., the training data) where the corresponding feature (component) appears, DisGeNET Disease: the disease in the DisGeNET database (i.e., the training data) where the corresponding feature (component) appears.

**Table S13.** The list of predicted Gv features above the corresponding Youden’s index for CKD following the removal of those in the training data, Related to Figure 3.

Database: modality, key_c: UMLS ID of the corresponding disease (i.e., the chronic kidney disease), Disease: the chronic kidney disease, component: the name of the predicted feature, keg component id: KEGG ID number for the predicted feature (component), obs: 0 as they are absent in the training data, sample count: count number in the output, freq: frequency in the output, roc_cutoff: the threshold cut-off value based on the Youden’s index, DisGeNET Database: the modality in the DisGeNET database (i.e., the training data) where the corresponding feature (component) appears, DisGeNET Disease: the disease in the DisGeNET database (i.e., the training data) where the corresponding feature (component) appears.

**Table S14.** The list of predicted Ae features above the corresponding Youden’s index for MI following the removal of those in the training data, Related to Figure 3.

Database: modality, key_c: UMLS ID of the corresponding disease (i.e., the myocardial infarction), Disease: the chronic kidney disease, component: the name of the predicted feature, keg component id: KEGG ID number for the predicted feature (component), obs: 0 as they are absent in the training data, sample count: count number in the output, freq: frequency in the output, roc_cutoff: the threshold cut-off value based on the Youden’s index, DisGeNET Database: the modality in the DisGeNET database (i.e., the training data) where the corresponding feature (component) appears, DisGeNET Disease: the disease in the DisGeNET database (i.e., the training data) where the corresponding feature (component) appears.

**Table S15.** The list of predicted Bm features above the corresponding Youden’s index for MI following the removal of those in the training data, Related to Figure 3.

Database: modality, key_c: UMLS ID of the corresponding disease (i.e., the myocardial infarction), Disease: the chronic kidney disease, component: the name of the predicted feature, keg component id: KEGG ID number for the predicted feature (component), obs: 0 as they are absent in the training data, sample count: count number in the output, freq: frequency in the output, roc_cutoff: the threshold cut-off value based on the Youden’s index, DisGeNET Database: the modality in the DisGeNET database (i.e., the training data) where the corresponding feature (component) appears, DisGeNET Disease: the disease in the DisGeNET database (i.e., the training data) where the corresponding feature (component) appears.

**Table S16.** The list of predicted Gv features above the corresponding Youden’s index for MI following the removal of those in the training data, Related to Figure 3.

Database: modality, key_c: UMLS ID of the corresponding disease (i.e., the myocardial infarction), Disease: the chronic kidney disease, component: the name of the predicted feature, keg component id: KEGG ID number for the predicted feature (component), obs: 0 as they are absent in the training data, sample count: count number in the output, freq: frequency in the output, roc_cutoff: the threshold cut-off value based on the Youden’s index, DisGeNET Database: the modality in the DisGeNET database (i.e., the training data) where the corresponding feature (component) appears, DisGeNET Disease: the disease in the DisGeNET database (i.e., the training data) where the corresponding feature (component) appears.

**Table S17.** The final list of the 30 predicted targets proteins for CKD, Related to Figure 3.

They are indicated by the gene/protein symbol (Symbol).

**Table S18.** The final list of the 37 predicted targets proteins for MI, Related to Figure 3.

They are indicated by the gene/protein symbol (Symbol).

**Table S19.** The raw data of Figure 4.

Each disease is indicated by UMLS ID (key_c) when available (otherwise, it is the same as the disease name). Each is indicated for CKD or MI (CKD_MI) and how many of the predicted genes for the corresponding disease appears in the training data for the corresponding disease (n_genes). The “n-genes” corresponds to “Count” in Figure 4.

**Table S20.** The raw data of Figure 5A.

The target is indicated as gene symbol (Symbol). The organ (Tissue) where the correspond target is expressed and its expression level (RNAExp) are shown.

**Table S21.** The raw data for Figure 5B.

The organ name (Tissue) and its p-value (pvalue), adjusted p-value (padj), and q-value (qvalue) are shown.

**Table S22.** The raw data for Figure 5C.

The target is indicated as gene symbol (Symbol). The cell-type (SCT) where the correspond target is expressed and its expression level (RNAExp) are shown.

**Table S23.** The raw data for Figure 5D.

The cell-type name (SCT) and its p-value (pvalue), adjusted p-value (padj), and q-value (qvalue) are shown.

**Table S24.** The raw data for Figure 6.

Those with padj<0.05 are listed.

**Table S25.** The KEGG pathway enrichment result for the CKD targets.

Those with p.adjust<0.5 are listed.

**Table S26.** The GO BP enrichment result for the CKD targets.

Those with p.adjust<0.5 are listed.

**Table S27.** The GO CC enrichment result for the CKD targets.

Those with p.adjust<0.5 are listed.

**Table S28.** The GO MF enrichment result for the CKD targets.

Those with p.adjust<0.5 are listed.

**Table S29.** The holdout validation of the therapeutic-indication (TI) prediction by the edit-distance based classifier (Method A), Related to Figure 7.

**Table S30.** The holdout validation of the side-effect (TI) prediction by the edit-distance based classifier (Method B), Related to Figure 7.

**Table S31.** The raw data (CKD, Method A) of Figure 8.

TI: the target disease (CKD), KEGG component_id: KEGG ID of the corresponding target, description: the name of the target, symbol: the symbol of the target, prob: probability of the prediction of the corresponding target, pathway: whether the target is in the same KEGG pathway as the clinically approved therapeutic target in the training data (1: in the same pathway, 0: in a different pathway)

**Table S32.** The raw data (CKD, Method B) of Figure 8.

TI: the target disease (CKD), KEGG component_id: KEGG ID of the corresponding target, description: the name of the target, symbol: the symbol of the target, prob: probability of the prediction of the corresponding target, pathway: whether the target is in the same KEGG pathway as the clinically approved therapeutic target in the training data (1: in the same pathway, 0: in a different pathway)

**Table S33.** The raw data (MI, Method A) of Figure 8.

TI: the target disease (CKD), KEGG component_id: KEGG ID of the corresponding target, description: the name of the target, symbol: the symbol of the target, prob: probability of the prediction of the corresponding target, pathway: whether the target is in the same KEGG pathway as the clinically approved therapeutic target in the training data (1: in the same pathway, 0: in a different pathway)

**Table S34.** The raw data (MI, Method B) of Figure 8.

TI: the target disease (CKD), KEGG component_id: KEGG ID of the corresponding target, description: the name of the target, symbol: the symbol of the target, prob: probability of the prediction of the corresponding target, pathway: whether the target is in the same KEGG pathway as the clinically approved therapeutic target in the training data (1: in the same pathway, 0: in a different pathway)

**Table S35.** The holdout validation of the side-effect (SE) prediction by the edit-distance based classifier (Methods A and B), Related to Figure 9.

**Table S36.** The raw data of Figure 9A.

**Table S37.** The raw data of Figure 9B.

**Table S38.** The raw data of Table 1.

The KEGG component IDs of the targets, the KEGG pathway IDs to which the targets belong to, GO IDs of the targets are also added to this table.

## References

Asico, L., Zhang, X., Jiang, J., Cabrera, D., Escano, C.S., Sibley, D.R., Wang, X., Yang, Y., Mannon, R., Jones, J.E., et al. (2011). Lack of renal dopamine D5 receptors promotes hypertension. Journal of the American Society of Nephrology: JASN 22, 82–89. 10.1681/ASN.2010050533.

Barabasi, A.L., Gulbahce, N., and Loscalzo, J. (2011). Network medicine: a network-based approach to human disease. Nature reviews. Genetics 12, 56–68. 10.1038/nrg2918.

Blei, D.M., Ng, A.Y., and Jordan, M.I. (2003). Latent dirichlet allocation. Journal of Machine Learning Research 3, 993–1022.

Buganim, Y., Faddah, D.A., Cheng, A.W., Itskovich, E., Markoulaki, S., Ganz, K., Klemm, S.L., van Oudenaarden, A., and Jaenisch, R. (2012). Single-cell expression analyses during cellular reprogramming reveal an early stochastic and a late hierarchic phase. Cell 150, 1209–1222. 10.1016/j.cell.2012.08.023.

Clemmer, J.S., Shafi, T., and Obi, Y. (2022). Physiological Mechanisms of Hypertension and Cardiovascular Disease in End-Stage Kidney Disease. Curr Hypertens Rep. 10.1007/s11906-022-01203-7.

De Hert, M., Detraux, J., and Vancampfort, D. (2018). The intriguing relationship between coronary heart disease and mental disorders. Dialogues Clin Neurosci 20, 31–40.

Garcia Del Valle, E.P., Lagunes Garcia, G., Prieto Santamaria, L., Zanin, M., Menasalvas Ruiz, E., and Rodriguez-Gonzalez, A. (2021). DisMaNET: A network-based tool to cross map disease vocabularies. Comput Methods Programs Biomed 207, 106233. 10.1016/j.cmpb.2021.106233.

Goh, K.I., Cusick, M.E., Valle, D., Childs, B., Vidal, M., and Barabasi, A.L. (2007). The human disease network. Proceedings of the National Academy of Sciences of the United States of America 104, 8685–8690. 10.1073/pnas.0701361104.

Hanrahan, J.P., de Boer, I.H., Bakris, G.L., Wilson, P.J., Wakefield, J.D., Seferovic, J.P., Chickering, J.G., Chien, Y.T., Carlson, K., Cressman, M.D., et al. (2020). Effects of the Soluble Guanylate Cyclase Stimulator Praliciguat in Diabetic Kidney Disease: A Randomized Placebo-Controlled Clinical Trial. Clin J Am Soc Nephrol 16, 59–69. 10.2215/CJN.08410520.

Hasin, Y., Seldin, M., and Lusis, A. (2017). Multi-omics approaches to disease. Genome biology 18, 83. 10.1186/s13059-017-1215-1.

Imai, J., Katagiri, H., Yamada, T., Ishigaki, Y., Suzuki, T., Kudo, H., Uno, K., Hasegawa, Y., Gao, J., Kaneko, K., et al. (2008). Regulation of pancreatic beta cell mass by neuronal signals from the liver. Science 322, 1250–1254. 10.1126/science.1163971.

Janssens, Y., Nielandt, J., Bronselaer, A., Debunne, N., Verbeke, F., Wynendaele, E., Van Immerseel, F., Vandewynckel, Y.P., De Tre, G., and De Spiegeleer, B. (2018). Disbiome database: linking the microbiome to disease. BMC Microbiol 18, 50. 10.1186/s12866-018-1197-5.

Ke, G., Meng, Q., Finley, T., Wang, T., Chen, W., Ma, W., Ye, Q., and Liu, T.-Y. (2017). LightGBM: A Highly Efficient Gradient Boosting Decision Tree.

Kho, J., Tian, X., Wong, W.T., Bertin, T., Jiang, M.M., Chen, S., Jin, Z., Shchelochkov, O.A., Burrage, L.C., Reddy, A.K., et al. (2018). Argininosuccinate Lyase Deficiency Causes an Endothelial-Dependent Form of Hypertension. American journal of human genetics 103, 276–287. 10.1016/j.ajhg.2018.07.008.

Koelwyn, G.J., Newman, A.A.C., Afonso, M.S., van Solingen, C., Corr, E.M., Brown, E.J., Albers, K.B., Yamaguchi, N., Narke, D., Schlegel, M., et al. (2020). Myocardial infarction accelerates breast cancer via innate immune reprogramming. Nature medicine 26, 1452–1458. 10.1038/s41591-020-0964-7.

Ku, E., Lee, B.J., Wei, J., and Weir, M.R. (2019). Hypertension in CKD: Core Curriculum 2019. Am J Kidney Dis 74, 120–131. 10.1053/j.ajkd.2018.12.044.

Lamy, J.B., Venot, A., and Duclos, C. (2015). PyMedTermino: an open-source generic API for advanced terminology services. Stud Health Technol Inform 210, 924–928.

Lloyd, G.G. (1987). Myocardial infarction and mental illness: a review. Journal of the Royal Society of Medicine 80, 101–104.

Menche, J., Sharma, A., Kitsak, M., Ghiassian, S.D., Vidal, M., Loscalzo, J., and Barabasi, A.L. (2015). Disease networks. Uncovering disease-disease relationships through the incomplete interactome. Science 347, 1257601. 10.1126/science.1257601.

Navarro, G. (2001). A guided tour to approximate string matching. ACM Computer Survey 33, 31–88. https://doi.org/10.1145/375360.375365.

Perakakis, N., Yazdani, A., Karniadakis, G.E., and Mantzoros, C. (2018). Omics, big data and machine learning as tools to propel understanding of biological mechanisms and to discover novel diagnostics and therapeutics. Metabolism 87, A1–A9. 10.1016/j.metabol.2018.08.002.

Reel, P.S., Reel, S., Pearson, E., Trucco, E., and Jefferson, E. (2021). Using machine learning approaches for multi-omics data analysis: A review. Biotechnol Adv 49, 107739. 10.1016/j.biotechadv.2021.107739.

Roller, S., and Schulte im Walde, S. (2013). A Multimodal LDA Model integrating Textual, Cognitive and Visual Modalities. Proceedings of the 2013 Conference on Empirical Methods in Natural Language Processing, 1146–1157.

Shahbazian, H., and Rezaii, I. (2013). Diabetic kidney disease; review of the current knowledge. J Renal Inj Prev 2, 73–80. 10.12861/jrip.2013.24.

Sharifi-Rad, J., Rodrigues, C.F., Sharopov, F., Docea, A.O., Can Karaca, A., Sharifi-Rad, M., Kahveci Karincaoglu, D., Gulseren, G., Senol, E., Demircan, E., et al. (2020). Diet, Lifestyle and Cardiovascular Diseases: Linking Pathophysiology to Cardioprotective Effects of Natural Bioactive Compounds. Int J Environ Res Public Health 17. 10.3390/ijerph17072326.

Shindo, T., Doi, S., Nakashima, A., Sasaki, K., Arihiro, K., and Masaki, T. (2018). TGF-beta1 promotes expression of fibrosis-related genes through the induction of histone variant H3.3 and histone chaperone HIRA. Scientific reports 8, 14060. 10.1038/s41598-018-32518-8.

Simoes, E.S.A.C., Miranda, A.S., Rocha, N.P., and Teixeira, A.L. (2019). Neuropsychiatric Disorders in Chronic Kidney Disease. Front Pharmacol 10, 932. 10.3389/fphar.2019.00932.

Suva, M.L., Riggi, N., and Bernstein, B.E. (2013). Epigenetic reprogramming in cancer. Science 339, 1567–1570. 10.1126/science.1230184.

Thomas, G., Sehgal, A.R., Kashyap, S.R., Srinivas, T.R., Kirwan, J.P., and Navaneethan, S.D. (2011). Metabolic syndrome and kidney disease: a systematic review and meta-analysis. Clin J Am Soc Nephrol 6, 2364–2373. 10.2215/CJN.02180311.

Underwood, E. (2021). A sense of self. Science 372, 1142–1145. 10.1126/science.372.6547.1142.

Valle, F., Osella, M., and Caselle, M. (2022). Multiomics Topic Modeling for Breast Cancer Classification. Cancers (Basel) 14. 10.3390/cancers14051150.

Wang, X., Wang, L.T., and Yu, B. (2022). UBE2D1 and COX7C as Potential Biomarkers of Diabetes-Related Sepsis. Biomed Res Int 2022, 9463717. 10.1155/2022/9463717.

Ward, P.S., and Thompson, C.B. (2012). Metabolic reprogramming: a cancer hallmark even warburg did not anticipate. Cancer cell 21, 297–308. 10.1016/j.ccr.2012.02.014.

Wen, Z., Nair, P., Deng, C.Y., Lu, X.H., Moseley, E., George, N., Lindvall, C., and Li, Y. (2021). Mining heterogeneous clinical notes by multi-modal latent topic model. PloS one 16, e0249622. 10.1371/journal.pone.0249622.

Wong, G., Staplin, N., Emberson, J., Baigent, C., Turner, R., Chalmers, J., Zoungas, S., Pollock, C., Cooper, B., Harris, D., et al. (2016). Chronic kidney disease and the risk of cancer: an individual patient data metaanalysis of 32,057 participants from six prospective studies. BMC Cancer 16, 488. 10.1186/s12885-016-2532-6.

Xing, Q.R., El Farran, C.A., Gautam, P., Chuah, Y.S., Warrier, T., Toh, C.D., Kang, N.Y., Sugii, S., Chang, Y.T., Xu, J., et al. (2020). Diversification of reprogramming trajectories revealed by parallel single-cell transcriptome and chromatin accessibility sequencing. Sci Adv 6. 10.1126/sciadv.aba1190.

Youden, W.J. (1950). Index for rating diagnostic tests. Cancer 3, 32–35. 10.1002/1097-0142(1950)3:1<32::aid-cncr2820030106>3.0.co;2-3.

Zhan, Q., Lv, Z., Tang, Q., Huang, L., Chen, X., Yang, M., Lan, L., and Shan, Q. (2021). Glycogen storage disease type VI with a novel PYGL mutation: Two case reports and literature review. Medicine (Baltimore) 100, e25520. 10.1097/MD.0000000000025520.

Zhang, S., Yamada, S., Park, S., Klepinin, A., Kaambre, T., Terzic, A., and Dzeja, P. (2021). Adenylate kinase AK2 isoform integral in embryo and adult heart homeostasis. Biochemical and biophysical research communications 546, 59–64. 10.1016/j.bbrc.2021.01.097.

Zheng, Y., Zhang, Y., and Larochelle, H. (2014). Topic Modeling of Multimodal Data: An Autoregressive Approach. 2014 IEEE Conference on Computer Vision and Pattern Recognition, 1370–1377. 10.1109/CVPR.2014.178.

